# Inflammation-like environments limit the loss of quorum sensing in *Pseudomonas aeruginosa*

**DOI:** 10.1101/2024.12.18.629113

**Authors:** Taoran Fu, Rosanna C.T. Wright, Danna R. Gifford, Christopher G. Knight, Michael A. Brockhurst

**Affiliations:** Division of Evolution, Infection and Genomic Sciences, School of Biological Sciences, Faculty of Biology, Medicine and Health, The University of Manchester, Manchester M13 9PT, UK; Department of Earth and Environmental Sciences, School of Natural Sciences, Faculty of Science and Engineering, The University of Manchester, Manchester M13 9PT, UK

**Keywords:** experimental evolution | quorum sensing | chronic infection | pathogen evolution | cystic fibrosis | *Pseudomonas aeruginosa*

## Abstract

Within-host environments are complex and multidimensional, making it challenging to link evolutionary responses of colonizing pathogens to causal selective drivers. Loss of quorum sensing (QS) via mutation of the master regulator, *lasR*, is a common adaptation of *Pseudomonas aeruginosa* during chronic infections. Here, we use experimental evolution in host-mimicking media to show that loss of QS is constrained by environmental factors associated with host inflammation. Specifically, environments combining oxidative stress and abundant free amino acids limited loss of QS, whereas QS loss was rapid in the absence of oxidative stress regardless of amino acids. Under oxidative stress, *lasR* mutations were contingent upon first decoupling regulation of oxidative stress responses from QS via mutations in the promoter region of the primary catalase, *katA*, or in the oxidative stress regulator, *oxyR*, enabling maintenance of oxidative stress tolerance. Together, our findings suggest that host inflammatory responses likely delay the loss of QS whilst colonizers undergo stepwise evolution, first adapting to survive lethal stressors before responding to other nutritional selective drivers that favour loss of QS.

**Significance Statement:** *Pseudomonas aeruginosa* is a common cause of chronic infections characterized by persistent inflammation. Host inflammatory responses alter within-host environments, including by increasing levels of antimicrobial stressors and releasing free amino acids through proteolysis. Here, we show stepwise adaptation of experimental *P. aeruginosa* populations to inflammation-like environments, first adapting to survive lethal stress by decoupling oxidative stress responses from quorum sensing, before then adapting to the nutritional conditions, delaying the loss of quorum sensing. These results highlight the power of using laboratory evolution experiments to disentangle the multidimensional selective forces driving pathogen adaptation in complex within-host environments.

## Introduction

Within-host environments are multidimensional, and this complexity is a barrier to understanding how pathogen adaptations are linked to the selective forces driving their evolution. This causality gap, in turn, limits our ability to create interventions that direct pathogen evolution towards desired outcomes, like reduced virulence. Even for well-studied pathosystems, such as *Pseudomonas aeruginosa* lung infections of cystic fibrosis (CF) patients, while we have detailed knowledge of the common pathways of genomic evolution (1), understanding of the causal selective drivers of particular adaptations is limited. Evolution experiments, directly observing microbial evolution in real time under strictly controlled conditions can be used to disentangle complex within-host selective environments (2). In particular, host-mimicking media is a powerful tool for hypothesis testing (3, 4), wherein experimental manipulations of key selective drivers expected to vary in hosts can be made and evolutionary responses tracked in real time (5–10). For example, addition of mucin drove increased population diversification (6, 9, 10), biofilm formation (9) and phage resistance (7), but weakened interaction between *P. aeruginosa* and the surrounding microflora (6). Oxygen availability and polyamines altered the evolution of phage resistance (8). Several studies have shown that nutrient and physiochemical conditions more strongly resembling CF lung sputum drive distinct evolutionary responses by *P. aeruginosa* (5, 8–11). For instance, creating CF-like levels of free amino acids, ionic profile and carbon sources drove the emergence of *mexT* mutations, associated with antibiotic resistance and quorum sensing, that were not observed in minimal glucose medium (11). CF-specific factors including bile salts, higher concentrations of mucin, DNA, free amino acids, sugars and host-derived antimicrobials drove greater antimicrobial resistance in media mimicking sinus or lung (8).

In this study, we focused on understanding the evolutionary impact of changes to the within-host environment driven by host inflammation. Persistent inflammation of the airway is common in both CF and non-CF bronchiectasis patients (3, 12, 13). The strength of inflammatory responses varies between individuals (14, 15), likely affecting the nature and strength of environmental selection experienced by colonising pathogens. During chronic airway infections, the host’s inflammatory response drives release of diverse effectors including antimicrobial compounds and enzymes, which drive changes in environmental conditions (16–18). Antimicrobials released by inflammatory immune cells include reactive oxygen and nitrogen species, as well as a cocktail of antimicrobial peptides (18). Neutrophilic proteolysis at infection sites by proteinases, neutrophil elastase and cathepsin G, targets and digests bacterial secreted proteins (19). As a consequence of host proteolysis, including degradation of host proteins, free amino acids are released, thus increasing their availability in inflamed airway microenvironments (3, 19). Consequently, colonising bacterial pathogens, such as *P. aeruginosa*, are exposed not only to antimicrobial stressors, like oxidative stress, but also to high levels of free amino acids liberated from proteins by the action of host proteases. How these nutritional and stressor axes of environmental selection interact to shape the trajectory of pathogen evolution is currently unclear.

A common *P. aeruginosa* adaptation observed in CF airway chronic infections is loss of acyl homoserine lactone (AHL) quorum sensing (QS) (20–23). Clinical isolates commonly exhibit loss of function mutations in the global regulator of AHL QS, *lasR*, which accumulate over time, particularly in more sustained and severe CF infections (24, 20, 22, 23). AHL QS positively regulates expression of a range of secreted products including extracellular proteases, including LasA, LasB, and alkaline protease (25, 26), catalase (27), siderophores (25, 28) and toxins (28–31). Bacterial proteases undergo a complex interaction with the host’s immune system in human infection (32–35), although the precise effect of losing protease secretion on infection severity and outcome is unclear (32–35). Loss of AHL QS is likely to increase susceptibility to host stressors, notably oxidative stress, because the expression of the primary catalase, KatA, and superoxide dismutases are positively regulated by LasR (27). Loss of quorum sensing, therefore, causes loss of expression of a suite of phenotypes many of which are linked to persistence and virulence. In addition, *lasR* mutations are also known to affect bacterial nutrient preferences. Metabolic studies show that *lasR* mutants more effectively utilize L- phenylalanine, L-lactic acid, L-arginine, and benefit from cross-feeding on citrate in co-culture with wild-type strains (36, 37). Moreover, *lasR* mutants can utilize amino acids released by neighbouring wild-type bacteria, without the cost of producing proteases, improving their competitive fitness in casein sole-carbon medium (38). Given the wide range of phenotypic effects caused by loss of QS, it is currently unclear how loss of QS is related to the environmental selective drivers present in the lung. Of particular interest, is to understand whether environmental changes driven by host inflammation, notably elevated levels of stressors and free amino acids, selection for *lasR* mutants given their association with severe disease.

Here, we experimentally evolved *P. aeruginosa* PA14 in environments mimicking airway-like environments with or without free amino acids and with or without oxidative stress in a fully factorial experimental design. Replicate populations were propagated by daily serial transfer for ∼ 250 generations. Population densities were tracked over time, alongside the frequency of colonies that had lost protease secretion, a proxy for loss of AHL QS. We used population sequencing and genetic analyses to compare the genomic targets of selection between treatments. Finally, we used a monoculture and coculture growth experiments with an isogenic *lasR* knockout strain (*ΔlasR*) to quantify the fitness effects of losing quorum sensing in these environmental conditions. Inflammation-like environments limited the evolutionary loss of quorum sensing, with oxidative stress in particular delaying the invasion of *lasR* mutants, which under oxidative stress were contingent upon prior mutations in promoter regions and regulators of oxidative stress responsive functions.

## Results

### Phenotypic adaptation to inflammation-associated environmental factors

To test how the availability of free amino acids and oxidative stress affected the evolutionary response of *P. aeruginosa*, we experimentally evolved replicate populations in an airway-mimicking medium (Synthetic CF sputum Medium (3), SCFM) modified to reflect environmental axes associated with inflammation. Specifically, we altered the availability of free amino acids, by replacing single amino acid resources with equal amounts of either casein (Casein SCFM) or casamino acids (CAA SCFM), and exposure to oxidative stress (with or without 2 mM hydrogen peroxide, OS+/-) in a fully factorial experimental design. Population densities varied over time and between treatments (Figure S1A). At first, population densities among environments were similar, with those in casein environments being slightly higher (LMM, day 1, nutrient resources as explanatory variable, with OS term being dropped by AIC comparison; nutrient term: *F1,22* = 8.83, *p* = 0.007; change in CFUs/ml = −0.46 × 10^9^, *p* = 0.007). However, whereas population densities increased over time in both nutrient environments without oxidative stress, such increases were not observed in either of the OS environments by day 42 (LMM, day 42, nutrient term: *F1,21* = 1.10, *p* = 0.307; OS term: *F1,21* = 7.57, *p* = 0.012). Together, these results suggest that OS constrained increases in population size, that were observed in both nutrient environments without OS.

Protease-deficient mutants (PDMs), which are likely to have gained AHL QS mutations, arose in all treatments but had contrasting invasion dynamics (Figure S1B, GLMM, interaction term between time and selection environments: *χ^2^3* = 2750.4670, *p* < 0.0001). Without OS, PDMs rapidly invaded to fixation in Casein SCFM OS−, whilst reaching only intermediate frequencies in most CAA SCFM OS− populations. OS delayed the emergence of PDMs in both nutrient conditions, doing so most strongly in CAA SCFM OS+ media (Figure S1B, survival analysis, in CAA SCFM, OS term coefficient = −2.6062, *z* = −2.35, *p* = 0.019). Protease deficiency was sometimes, but not always associated with a small colony variant phenotype (Figure S2). At the endpoint of the selection experiment, PDMs reached high frequency in most populations, except those in the CAA SCFM OS+ environment.

By the endpoint of the experiment, increased tolerance of hydrogen peroxide had evolved in all populations selected under OS regardless of the nutrient environment, whereas slightly reduced tolerance of hydrogen peroxide evolved in populations selected without OS (Figure S3, GLMM, count varied by interaction term between OS and MIC level, in CAA SCFM: *χ^2^1* = 20.96, *p* < 0.001; in Casein SCFM: *χ^2^1* = 19.86, *p* < 0.001). This pattern is consistent with previous work suggesting a trade-off governing short-term adaptation to environmental nutrients versus OS (39). In contrast, we observed no consistent change in resistance to the common clinical antibiotics ciprofloxacin or tobramycin associated with any of the treatments, defined as minimum inhibition concentration higher than ancestral PA14, although some individual populations showed enhanced resistance to both antibiotics (Figure S4).

### Oxidative stress drives genomic divergence and contrasting evolutionary dynamics

To understand the genomic responses to selection we obtained population whole genome sequences at days 14, 28 and 42. Across all evolved lines and time points, we observed 25 synonymous SNPs, 204 nonsynonymous SNPs and small indels, 81 mutations in intergenic regions and 3 larger structural variants (Figure S5 and Table S1, S2, S3, and S4). Because OS is mutagenic it could potentially have caused mutations, as such we first tested for mutational bias among treatments by analysing the distributions of synonymous mutations. However, the abundance of synonymous mutations did not significantly differ between selection environments (GLMM, at gene level, adding OS term: *χ^2^1* = 2.05, *p* = 0.152; adding free amino acids term: *χ^2^1* = 0.25, *p* = 0.616), but was instead dependent on the length of the gene and position on the genome (GLMM, at gene level, the length of the gene: *χ^2^1* = 156.23, *p* < 0.001; position on the genome: *χ^2^1* = 7.08, *p* = 0.008).

Focusing on the subset of nonsynonymous mutations and mutations in intergenic regions, we next tested whether the different selective factors had caused genomic divergence between treatments. Comparison of pairwise similarities of mutational profiles between replicate populations per sampled time point showed greater similarity within than between treatments at all time points (Permutation ANOVA of Jaccard index within versus between treatments: *F1,1722* = 134.912, *R^2^* = 0.07, *p* < 0.001; time: *F2,1722* = 39.71, *R^2^* = 0.04, *p* < 0.001; interaction: *F2,1722* = 2.08, *R^2^ =* 0.002, *p* = 0.081). This pattern of genomic divergence was driven by differing evolutionary paths being followed with versus without OS selection. Specifically, 10/12 replicate lines selected with OS gained mutations in the upstream region of *katA*, regardless of the nutrient conditions (Mann- Whitney U test, *W* = 24, *p* = 0.174). All 3 of the observed mutations occurred within the σ70 promoter region (as predicted by SAPPHIRE (40)), with a higher occupancy score for OxyR binding motifs (41, 42) indicating greater transcription factor binding efficiency, and thus would be likely to affect the transcription of *katA*. Moreover, 9/12 replicate lines selected with OS gained mutations in *oxyR* (*PA14_70560*) at least at one time point, with mutations in this gene being more commonly selected under free amino acid nutrient conditions (GLM, populations with *oxyR* mutation across treatments and time points, time dropped by AIC comparison, treatment: *χ^2^1* = 4.88, *p* = 0.027). Unlike OS selection, selection arising from the nutrient conditions did not appear to drive significant genomic divergence between treatments consistently across time points.

Although not consistent across both nutrient conditions, mutation in *imuB* (*PA14_55600*, SOS response DNA repair) also appeared to be associated with OS. In the Casein SCFM OS+ treatment, *imuB* mutation rose to high frequency between days 14 and 42 of the experiment, in 3/6 replicate populations. ImuB is a damage-inducible polymerase known to contribute to the SOS response and UV-induced mutagenesis DNA repair (43, 44) but its specific function in response to OS is less well understood.

The dynamics of mutations affecting *lasR* broadly tracked the dynamics of PDMs in most populations (Figure 1). Whilst nonsynonymous SNPs in *lasR* were observed in some populations, others acquired large deletions containing ∼ 30 genes from *PA14_45700* to *PA14_46100*, including QS system-related genes (*lasI*, *lasR* and *rsaL*), predominantly in the Casein SCFM treatment. These deletions affected several adjacent genes, including the flagellar operon (*fliL-fliM-fliN-fliO- fliP-fliQ-fliR-flhB-PA14_45710-PA14_45700*), a two-component regulatory system operon (*PA14_45880-PA14_45870*), a RND efflux operon (*PA14_45910, PA14_45890*), a putative cation- transporting P-type ATPase gene (*PA14_45970*), a putative acetyltransferase gene (*PA14_45980*), and an operon contributing to the encoding of a putative ABC transporter ATP-binding protein (*PA14_46010-PA14_46000*). Without OS, free amino acids accelerated the emergence of *lasR* mutants to detectable levels, with 3/6 appearing by day 14 in CAA SCFM, compared to none in Casein SCFM (Figure 1).

**Figure 1.**
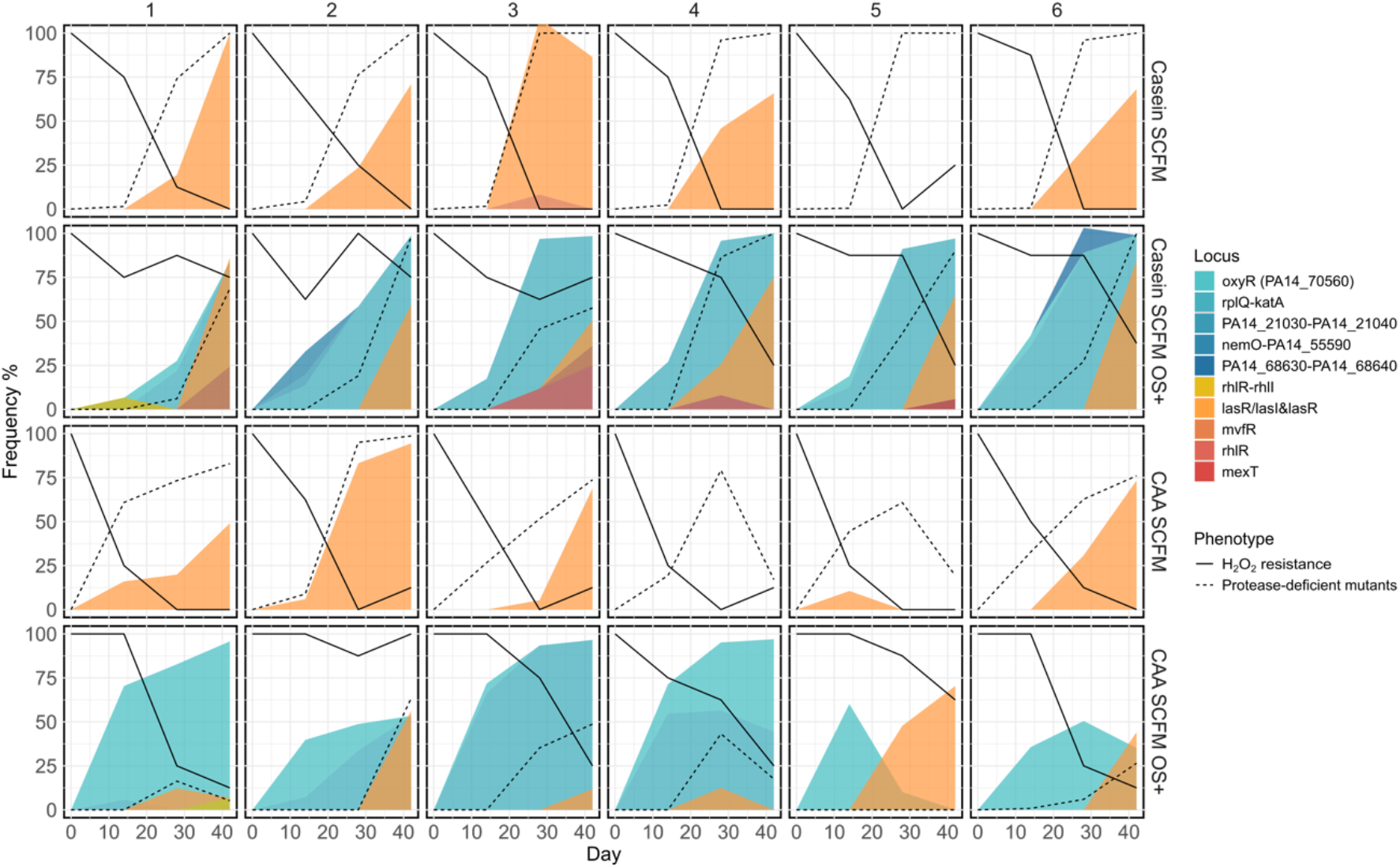
Frequencies of key phenotypes and mutations over time. Frequency of hydrogen peroxide resistance (n=8, solid lines), protease-deficient mutants (PDMs, a proxy for loss of AHL QS, detection limit at 10^6^ cfu, dashed lines) and mutations relevant to QS (orange) and oxidative stress (blue-green) responses on days 14, 28, and 42. Each plot represents an individual replicate population arranged in rows per treatment (see labels). Populations were serially transferred in medium mimicking CF sputum (SCFM) modified to reflect environmental axes associated with host inflammation. Specifically, we altered the availability of free amino acids by replacing single amino acid resources with the same amounts of casamino acids (CAA SCFM) or casein (Casein SCFM), and levels of oxidative stress, by culturing with or without addition of of 2 mM hydrogen peroxide (OS+/−)

However, although loss of AHL QS, was common across all treatments, it was delayed by OS (Figure 1, survival analysis, OS term coefficient = -2.06, *z* = -3.30, *p* < 0.001; nutrient conditions term coefficient = -0.56, *z* =-1.31, *p* = 0.190). Under OS, *lasR* mutations were consistently preceded by mutations either in the promoter region upstream of *katA* or in *oxyR* reaching high frequency. Populations with mutations in the *katA* promoter showed higher tolerance to hydrogen peroxide (Figure 1; GLMM, frequency of H2O2 resistant (0 ∼ 1) as response, *katA* promoter mutations on day 28 and 42; mutations in *katA* upstream: *χ^2^1* = 9.07, *p* = 0.003; time: *χ^2^1* = 1.59, *p* = 0.207; interaction: *χ^2^1* = 0.655, *p* = 0.418), and subsequent *lasR* mutations in these populations did not cause complete loss of hydrogen peroxide tolerance, unlike populations without *katA* promoter mutations. Together, the mutational and phenotypic dynamics suggest that under oxidative stress, *lasR* mutations were contingent upon prior adaptation to OS through *katA* promoter or *oxyR* mutations (GLMM frequency of *lasR* mutation in OS+ treatments: frequency of *katA* promoter or *oxyR* mutations at the prior time point: *χ^2^1* = 134.31, *p* < 0.001; time: *χ^2^1* = 0.04, *p* = 0.85; interaction: *χ^2^1* = 4.54, *p* = 0.033).

Mutations at several other loci could be linked to specific treatments or changes in phenotype. *psdR* mutations were only observed in Casein SCFM environments, consistent with its function as a regulator of dipeptide metabolism (45–47). In Casein SCFM OS− replicate 5, the invasion of PDMs was not associated with mutations affecting *lasR*, however mutations in 3 genes closely tracked PDM frequency, including a mutation affecting *PA14_66580* encoding a putative Type 2 secretion system protein, possibly involved in protease secretion. Although loss of flagellar motility occurred across all treatments (Figure S6), mutation of the flagellar transcriptional regulator, *fleQ*, was only observed in CAA SCFM, whereas mutations in other flagellum-associated genes occurred in other treatments.

### PA14*ΔlasR* is more susceptible to oxidative stress but beneficial in both nutrient environments

To test the effects of *lasR* loss-of-function, we compared the growth and competitive fitness of PA14 against an isogenic knock-out mutant, PA14*ΔlasR*, across the environmental conditions used in the selection experiment. PA14*ΔlasR* was much more susceptible to hydrogen peroxide than PA14 wild-type (Figure 2A), explaining why *lasR* mutants were unable to invade under OS without prior mutation of the *katA* promoter and/or *oxyR*. Consistent with the invasion of *lasR* mutants under both nutrient conditions in the selection experiment, PA14*ΔlasR* outcompeted wild-type PA14 in both nutrient environments (paired *t*-test, in Casein SCFM: *t5* = 4.90, *p* = 0.004; in CAA SCFM: *t5* = 3.15, *p* = 0.026), confirming that *lasR* mutants are fitter than the wild-type regardless of the availability of free amino acids (Figure 2B and S7). Notably, however, while co-culture with PA14 increased the abundance of PA14*ΔlasR* compared to monocultures in casein, the abundance of PA14*ΔlasR* was decreased relative to monocultures by co-culturing with PA14 in CAA (Figure 2B). This is consistent with previous findings that *lasR* mutants benefit from the protease activity of neighbouring wild-type cells liberating free amino acids through digestion of casein, but are also more efficient at utilizing various free amino acids than wild-type cells (36–38). Accordingly, over the entire 42-day experiment, we observed a positive correlation between population density and the frequency of PDMs only in the CAA SCFM environment (Figure 2C, LMM, PDM frequency: *χ^2^1* = 102.58, *p* < 0.001), suggesting that PDMs improve population-level adaptation only in the absence of oxidative stress and when amino acids are freely available.

**Figure 2.**
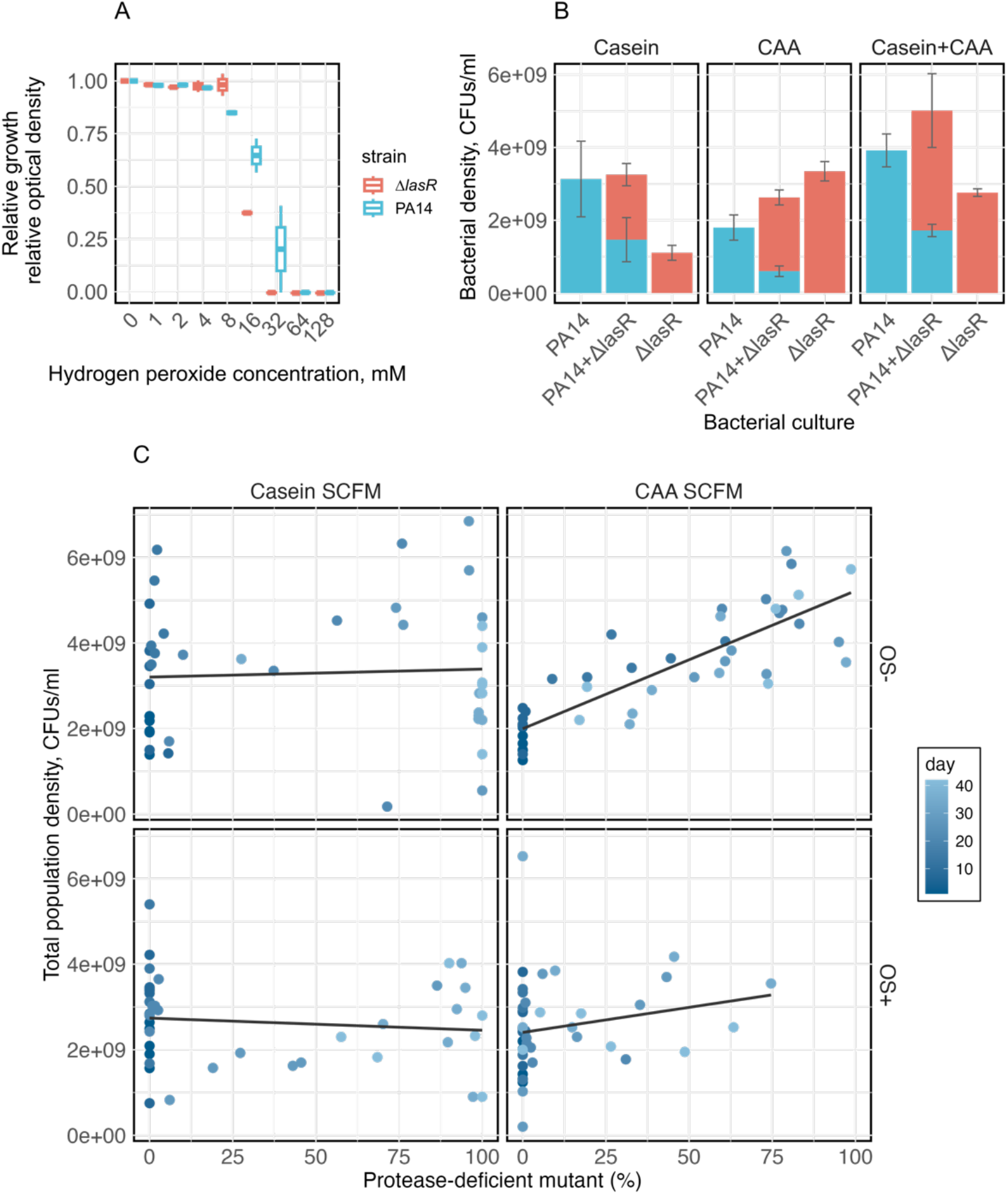
Effects of loss of QS on monoculture and coculture growth and population densities. (A) Relative optical densities of wild-type PA14 (blue-green) and PA14 *ΔlasR* (red) at exposure to different concentrations of hydrogen peroxide, relative to stress-free, after 24 hours, show that PA14 *ΔlasR* is more susceptible to hydrogen peroxide than wild-type PA14. (B) Endpoint population density of PA14 (blue-green) and PA14 *ΔlasR* (red) in mono-culture and co-culture in stress-free Casein, CAA SCFM or a mixture of both after 24 hours. Co-culturing with wild-type PA14 affects bacterial density that differs in Casein SCFM versus CAA SCFM. (C) The total population density positively correlates with the frequency of evolved protease-deficient mutants (PDMs) only in stress-free CAA SCFM. Color gradient indicates the day of observation.

## Discussion

*P. aeruginosa* undergoes a suite of characteristic adaptations during human chronic airway infections, including loss of QS (48–51). Here, we show that evolutionary loss of QS is limited by inflammation-like environments. Whereas QS was rapidly lost under both nutrient conditions in the absence of oxidative stress due to higher fitness of *lasR* mutants, loss of QS was constrained by a combination of free amino acids and oxidative stress, two environmental factors associated with inflammation. Population sequencing revealed that, under oxidative stress, *lasR* mutations were contingent upon first acquiring mutations in the *katA* promoter region or *oxyR*. These mutations were associated with maintaining high tolerance to hydrogen peroxide, enabling *lasR* mutations, which otherwise caused increased sensitivity to hydrogen peroxide. Together, our results suggest that host inflammatory responses are likely to delay the evolutionary loss of quorum sensing in *P. aeruginosa* chronic infections, and as such the correlation between QS loss and severe disease (23, 32, 52–55) is unlikely to be driven by the inflammatory response *per se*.

AHL QS coordinates the expression of a wide range of important bacterial traits (26, 56–58). While such integrated regulatory systems clearly provide adaptive benefits in an organism’s natural niche, individual regulated traits may experience contrasting selection pressures when exposed to novel environments, such as the human airway (48, 59–62). Such potential conflicts among co-regulated traits are exemplified here by the contrasting effects of stressor versus nutritional selection pressures upon *lasR* mutants. Selection for the loss of *lasR* mediated by the nutritional environments required first the decoupling of regulation of oxidative stress responses from *lasR* in environments with hydrogen peroxide. This regulatory decoupling arose by two alternate mechanisms, either promoter mutations upstream of the primary catalase, *katA*, or mutation of the primary hydrogen peroxide transcriptional activator gene, *oxyR*, permitting bacteria to maintain high OS tolerance following subsequent mutation of *lasR*. Both LasR and OxyR have been shown to upregulate the expression of *katA* in response to hydrogen peroxide (27, 63), and the *katA* promoter mutations we observed are predicted to enhance OxyR binding. As such our data suggest a model whereby, under OS, reduced LasR-mediated regulation of *katA* evolved whilst *katA* expression became predominantly driven by OxyR.

Release of reactive oxygen species is a common host defence mechanism, including as part of the human inflammatory response, and is increasingly recognised as a key selective pressure shaping pathogen evolution (19, 64, 65). OS has a range of potential effects on bacteria, including elevating the mutation rate (66), inducing stress responses (64), and eliciting antibiotic resistance (64). Here, we used physiologically relevant levels of oxidative stress to mimic OS responses in the host lung environment but observed no differences in the number of synonymous mutations with or without OS, suggesting that the mutagenic effects of OS may not be important within the CF lung. Furthermore, we observed no differences among treatments in collateral evolution of antibiotic resistance, again suggesting that at physiological levels, OS is unlikely to be a major driver of antibiotic resistance, consistent with a clinical study (49). By contrast, OS had a strong ecological effect, suppressing population densities over the duration of the experiment compared to treatments without OS, and was the main driver of genomic divergence between treatments via selection upon OS responsive genes. Together with the findings in a previous study (39), the result suggests that OS and how this interacts with population ecology at the site of infection is a key selective force in within-host environments. Indeed, our results suggest a potentially general rule of adaptation in complex multidimensional selection environments whereby organisms must first adapt to stressors to avoid extinction, before adapting to other environmental axes, such as nutritional niches.

Our experiments used a host-mimicking medium (3) supplying a physiologically-relevant amount of carbohydrate alongside amino acids as either free amino acids or as proteins. *lasR* mutants were favoured in both nutritional environments. In free amino acid environments, this likely arises from the higher metabolic efficiency of *lasR* mutants compared to wild-type cells for various abundant amino acids (36). Monocultures of PA14*ΔlasR* thus reached higher population densities in CAA than either PA14 or cocultures, and in the selection experiment we saw a positive correlation between population density and the frequency of PDMs (a proxy for *lasR* mutation) in CAA SCFM OS−. Thus, in free amino acid environments loss of AHL QS is individually beneficial. While extracellular proteases can play a crucial role in bacterial nutrient acquisition (67–69), ecological competition (67, 69), and pathogenicity (69, 70), they are costly to produce and susceptible to exploitation by nonproducing cheaters who benefit from the resources liberated whilst paying no cost (38, 71). As such, in casein environments, where amino acids are supplied as proteins, *lasR* mutants gain an additional indirect fitness benefit by exploiting the proteolysis performed by neighbouring wild-type cells, consistent with previous studies using casein sole-carbon media environments (72, 73).

Overall, we show that inflammation-like environments limit the evolutionary loss of QS by delaying the invasion of *lasR* mutants. This is because, although *lasR* mutants are fitter than wild-type in environments with high levels of amino acids, they are selected against by OS. For *lasR* mutants to invade in inflammation-like conditions, bacteria must first adapt to OS by decoupling the regulation of OS responses from QS, exemplifying the probable primacy of stressor adaptation over nutritional adaptation in multidimensional selective environments.

## Materials and Methods

### Bacteria and culture condition

*P. aeruginosa* UCBPP-PA14 (74) glycerol stock was streaked on King’s B (KB) agar plate and grown overnight. KB is a general non-selective medium for bacterial growth which contains 20g/L proteose peptone No.3, 10ml/L glycerol, 1.5g/L K2HPO4, and 1.5g/L MgSO4, supplemented with 15 g/L for agar plates. Overnight cultures or agar plates were incubated overnight at 37°C and overnight cultures were shaken at 180 rpm if not specified. If not otherwise indicated, bacteria were preconditioned to grow overnight in the same medium before assays.

### Evolution experiment

We evolved single clones of PA14 in a medium mimicking CF sputum (SCFM) (3) but modified to contain or not contain free amino acids by replacing single amino acid resources with the same amounts of casamino acids (CAA SCFM) or casein (Casein SCFM) and with or without a calculated final concentration of 2 mM hydrogen peroxide (OS+/−), the concentration defined from a previous study (39) to cause a sublethal effect of OS. Among them, Casein SCFM OS− represents the environment with least inflammation, Casein SCFM OS+ or CAA SCFM OS− represents either oxidative stress or availability of free amino acids, and CAA SCFM OS+ represents the highest level of inflamed environment with both factors. Additional growth curves were measured in advance to confirm growth had reached stationary phase after 24 hours in different media. Replicate populations were pre-adapted to the corresponding nutrient environment until reaching the stationary phase, before setting up the growth in one of four selective environments on day 1. Daily transfer was carried out by transferring 1% of the growth culture into fresh media, incubated at 37°C with shaking at 180 rpm for at least 22 hours. Each selective environments include 6 replicate populations. To track the emergence of evolved PDMs in the evolution experiment, we plated diluted culture on 3% skim milk plate on the experiment set-up day and then every seven days after that. The proportion of colonies without clearing rings against the total colony forming units (CFUs) was determined as frequency of PDMs, likely representing the frequency of loss of quorum sensing (22, 38). Eight colonies per population were picked randomly through a grid and cultured and frozen at -80°C every seven days. Meanwhile, 200 μl from each population was also frozen.

### Hydrogen peroxide susceptibility assay

To test for the hydrogen peroxide susceptibility of *lasR* mutant and evolved strains from day 42, we inoculated the overnight culture of strains in 96-well plates into fresh Muller Hinton broth 2 (MH2, Millipore, Merk, Germany) supplemented with a range of hydrogen peroxide at a calculated final concentration of: 0, 1, 2, 4, 8, 16, 32, 64, 128 mM. The optical density at 600 nm was measured after incubation for 24 hours at 37°C. The correct optical density was calculated by subtracting the reads of the clean medium from the raw reads. Growth inhibition was calculated by the decrease in the ratio of the correct optical density relevant to H2O2-free growth. A growth inhibition larger than 95% is considered negligible growth after 24 hours. The 96-well plates were left on the bench for an additional 48 hours to allow for slow growth, which helps distinguish between susceptible and resistant strains. For testing the frequency of low susceptible evolved strains in all populations, eight evolved strains isolated from each population on day 7, 14, 28, 35, and 42 were revived in MH2 and inoculated into fresh MH2 supplemented with the same concentration of hydrogen peroxide, about 64 mM. The growth inhibition relevant to full growth without hydrogen peroxide was calculated, with an inhibition over 95% regarded as a full inhibition of growth. Assay 96-well plates were left on the bench for 48 hours more and negligible growths were recorded.

### Pooled genomic sequencing

Samples of the ancestral PA14 clone and from each of the experimental evolution line populations from day 14, day 28, and day 42 were defrosted from frozen stocks. Serial dilutions of each sample were plated onto KB agar plates to a density of > 300 colonies per plate after overnight incubation. Colonies were washed from each agar plate with M9 and DNA was extracted using a DNeasy kit (Qiagen, Germany) as per the manufacturer’s protocol. Additional DNA purification was performed using magnetic bead kits for samples that yielded low DNA concentration or purity determined by gel electrophoresis, Qubit and nanodrop (beyond A260/A280 = ∼ 0.8 or 1.8 < A260/A230 < 2.2). Library preparation was performed by the Centre for Genomic Research Liverpool (CGR). Sequencing was performed by CGR using Illumina NovaSeq 2×150-bp paired-end reads at ∼ 1,150× read depth. Data QC were performed by CGR, where the raw Fastq files were trimmed for the presence of Illumina adapter sequences using Cutadapt v1.2.1 (75) and further trimmed using Sickle v1.200 (76) with a minimum window quality score of 20.

### Variant calling

Single nucleotide polymorphisms (SNPs) and small indels were called against the PA14 reference genome in NCBI (NC_008463.1) using a workflow adapted for *P. aeruginosa* pool sequencing, including alignment using bwa mem v0.7.17 (77), picard v3.1.0 (78) to remove duplicate reads, genomeCoverageBed from bedtools v2.31.0 (79) for coverage analysis, samtools v1.18 (80) to generate pileup outputs from bam files, varscan v2.4.6 (81) using mpileup2cns flag for variant calling at minimal coverage 5, minimal variant frequency 5%, with p-value 0.05. Variants were annotated using snpEff v5.2 (82) and relevant to PAO1 GO terms (83). Large structural variants were identified using delly v1.1.7 (84) and verified visually using IGV v2.17.0 (85) and bam files. Breseq v0.38.1 (86) was used as a complementary bioinformatic approach to assess robustness and consistency of variant calling using the bespoke pipeline. Ancestral PA14, with no considerable variant being observed compared to the PA14 reference genome in NCBI (NC_008463.1), served as a reference to filter out variants in the evolved populations. Variants with frequency > 5% were included in downstream analyses.

### Competition assay

To understand the fitness of loss of AHL quorum sensing relative to wild-type (PA14), we mono or co-culture both of PA14 and an isogenic *lasR* knockout strain (PA14Δ*lasR*) in Casein SCFM OS− and CAA SCFM OS−. Bacterial cultures were diluted and plated onto milk agar plates before and after incubating for 24 hours at 37°C to determine CFUs at stationary phase. Additional growth curves were measured to confirm reaching stationary phase after 24 hours of growth in both nutrient conditions. To investigate the fitness of PA14Δ*lasR* at a larger range of starting ratio, overnight cultures were washed using M9 solution and used to set up competition assay at a starting ratio varying from 1:100, 1:10, 1:1, 10:1 to 100:1, in triplicate.

### Antibiotic resistance

To investigate the evolution of antibiotic resistance in the evolution experiment, we determined minimal inhibitory concentration (MIC) of common clinical antibiotics of evolved strains, ciprofloxacin and tobramycin MICs. The range of ciprofloxacin and tobramycin concentration covered the clinical breakpoints. Evolved strains were recovered from frozen stock and inoculated into fresh MH2 supplemented with either ciprofloxacin at a concentration of 0, 0.125, 0.25, 0.5, 1, or 2 ug/ml or tobramycin at a concentration of 0, 0.5, 1, 2, 4, or 8 ug/ml.

### Swimming motility assay

To investigate the loss of flagellar motility in the evolution experiment, individual colonies were inoculated from the frozen stock, cultured overnight, calibrated to optical density OD=∼ 0.3 at 600 nm and stabbed into a 0.3% agar plate (1xM8 solution, 0.5% casamino acids, 0.2% glucose, and 1 mM MgSO4) (87). After incubation at 37 °C for 8 h, the swimming motility was determined by an obvious radial migration through the agar. PA14, a PA14 *lasR* knock-out strain, PAO1, and a PAO1 *fleQ* knockout strain served as controls.

## Statistical analysis

Statistical analysis and visualisation were conducted in R v4.3.1. Several packages were used for data preparation and visualisation including ‘tidyverse’ (88), ‘ggplot2’ (89) and ‘patchwork’ (90). Linear mixed effect models (LMM) were built using package ‘lme4’ (91) or ‘glmmTMB’ (92) for generalised linear mixed effect model (GLMM), Chi-square test through ‘car’ (93). To pick explaining terms for variation in population density on day 1 and 42, we used an LMM with lineage as the random effect, and dropped single term if AICs decreased in the simpler model. Survival analysis was used to estimate the timing of PDMs or *lasR* mutation emergence (0 or 1), with a positive emergence regarded as observed single PDM on milk agar plate (equivalent to about 10^7^ cfu/ml) or a detected *lasR* mutation, repectively. Statistical models, GLMM of negative binomial distribution, were used to explain variation in number of synonymous mutations by comparing all combinations of the length of the gene, position on the chromosome, time and availability of free amino acids and oxidative stress, and models with lowest AICs and significant Chi-square test results were picked. Generalised linear models (GLM) with a negative binomial distribution were used to analyze variation in the count of populations with mutations in genes of interest across treatments and time points. Beta-family GLMMs were used to analyze mutation frequencies of interest at the gene level, including tests for correlation between the frequency of *lasR* mutations across replicate populations and time points and the frequency of *katA* promotor regions or *oxyR* mutations at the previous time point, with the interaction by time. Parallel evolution or similarity between all possible pairs of sample, quantifying using the Jaccard index, the number of gene targeted in common divided by total number of loci targeted. Parallel evolution within same treatment versus between treatments was characterized by permutational ANOVA (1,000 permutations), using ‘adonis’ (‘vegan’ R package (94))

## Acknowledgments

We are grateful to E. Westra and T. Taylor for providing the knockout strains used in this study, A. Kottara for providing lab protocols, and A. Buckling and F. Whelan for valuable suggestions on the manuscript. This work was supported by funding from the Wellcome Trust to MAB (220243/Z/20/Z) and the BBSRC to R.C.T.W. and M.A.B. (BB/T014342/1).

## Author Contributions

T.F., R.C.T.W, D.R.G., C.G.K. and M.A.B designed research; T.F. performed experiments; T.F., R.C.T.W and M.A.B contributed analytic tools; T.F. and M.A.B analyzed data; T.F. and M.A.B. wrote the first draft of the manuscript; all authors reviewed and revised the final manuscript.

