## Supplementary figures for "Inflammation-like environments limit the loss of quorum sensing in *Pseudomonas aeruginosa*"

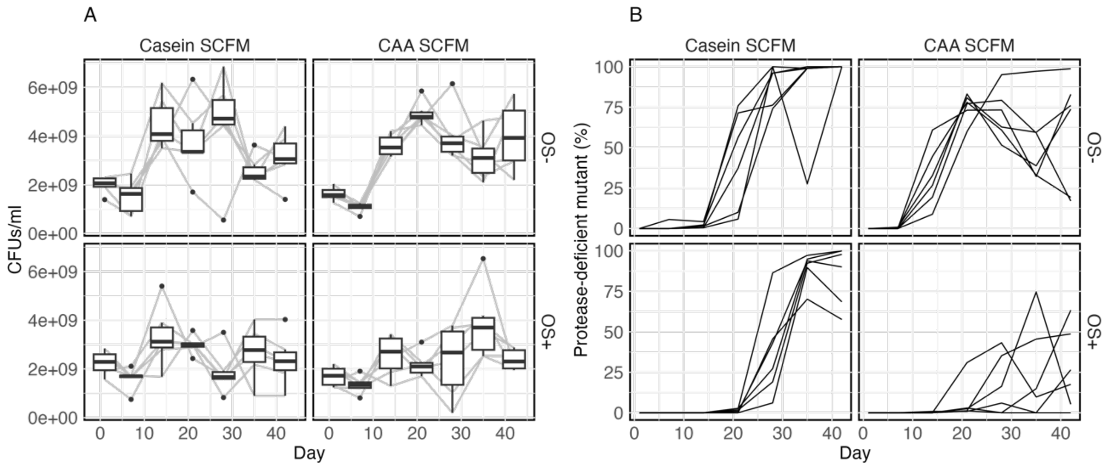


**Figure S1.** Population densities and frequencies of protease-deficient mutants over time. Differences in environmental factors associated with inflammation varied the population density (A) and the frequency of evolved protease-deficient mutants, PDMs (B) over time. Box plot tracks the average and overall distribution of population density at each detected time point within each selective environment. Each line shows the tracked population density or PDMs frequency along the daily passage in a single population, grouped by selective environments (Casein SCFM or casamino acids, labeled CAA SCFM, and with or without supplemented 2 mM hydrogen peroxide, OS+/-).

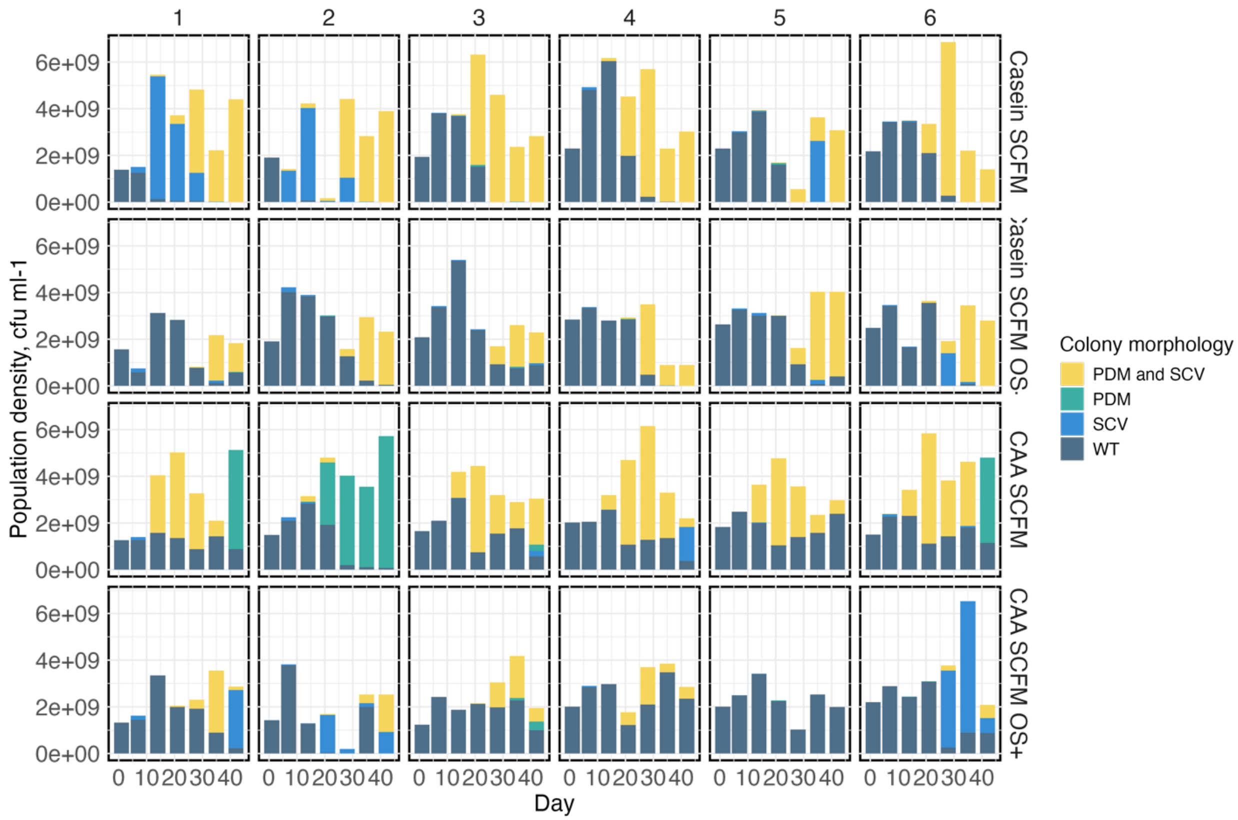


**Figure S2.** Distinct colony morphology types across treatments over time. Four classes of colony morphologies were observed: wild-type PA14-like (WT), protease-deficient mutant (PDM), small colony variant (SCV), and small colony variant with a protease-deficiency (PDM and SCV).

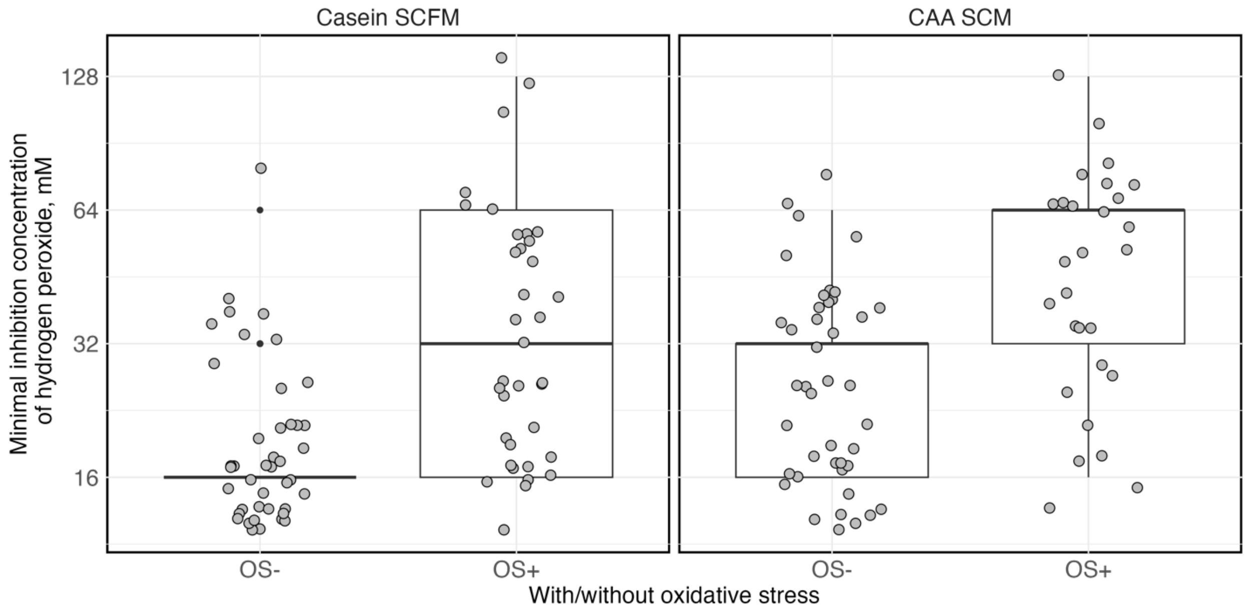


**Figure S3.** Evolution of reduced susceptibility to hydrogen peroxide following selection under oxidative stress. Hydrogen peroxide minimum inhibitory concentration of evolved clones taken from day 42 (*n*=8 per population, 6 populations per selective environments).

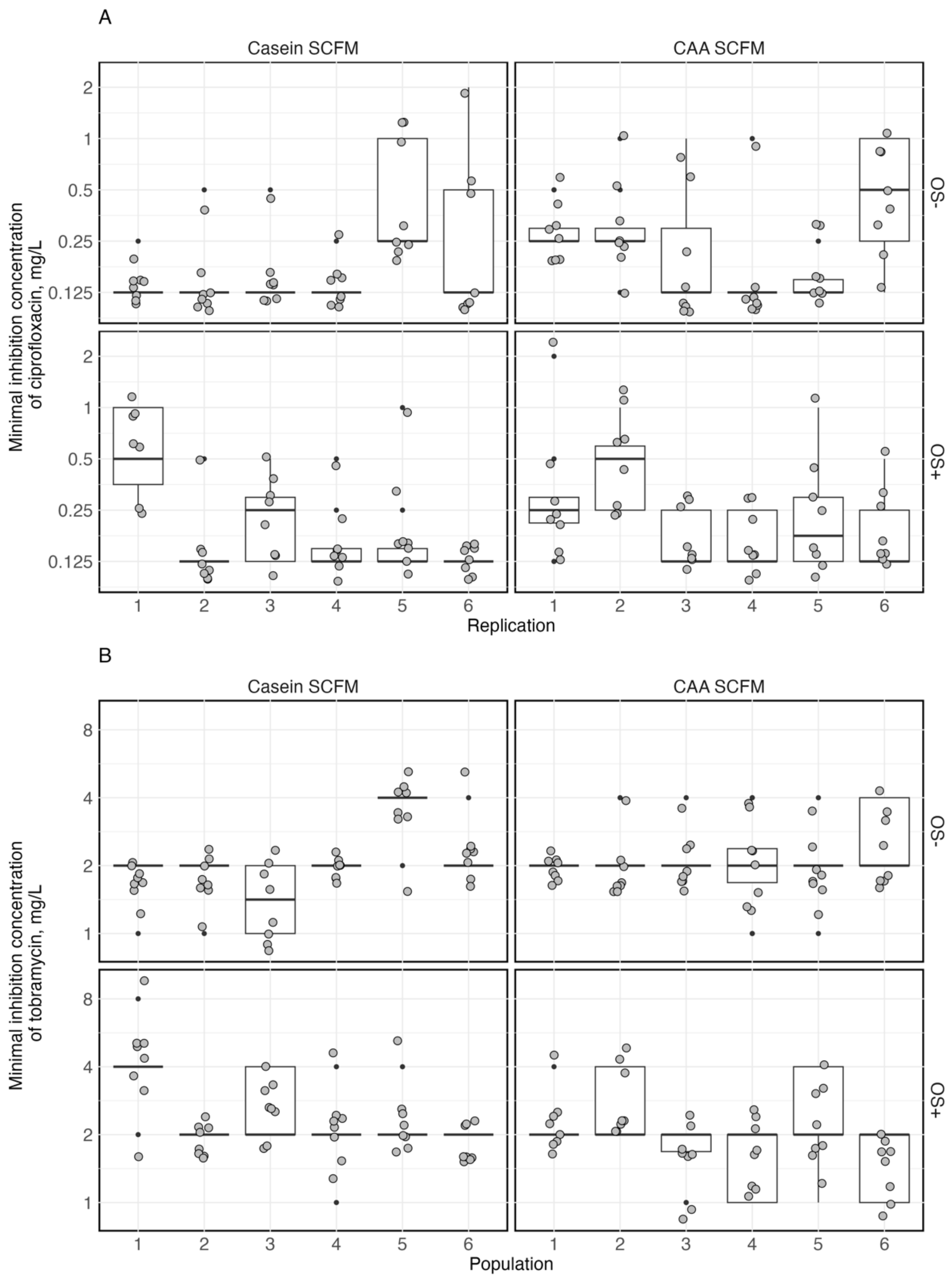


**Figure S4.** Minimum inhibitory concentration of ciprofloxacin (A) or tobramycin (B) of evolved clones taken from day 42 ($n$=8 per population, 6 populations per selective environment).

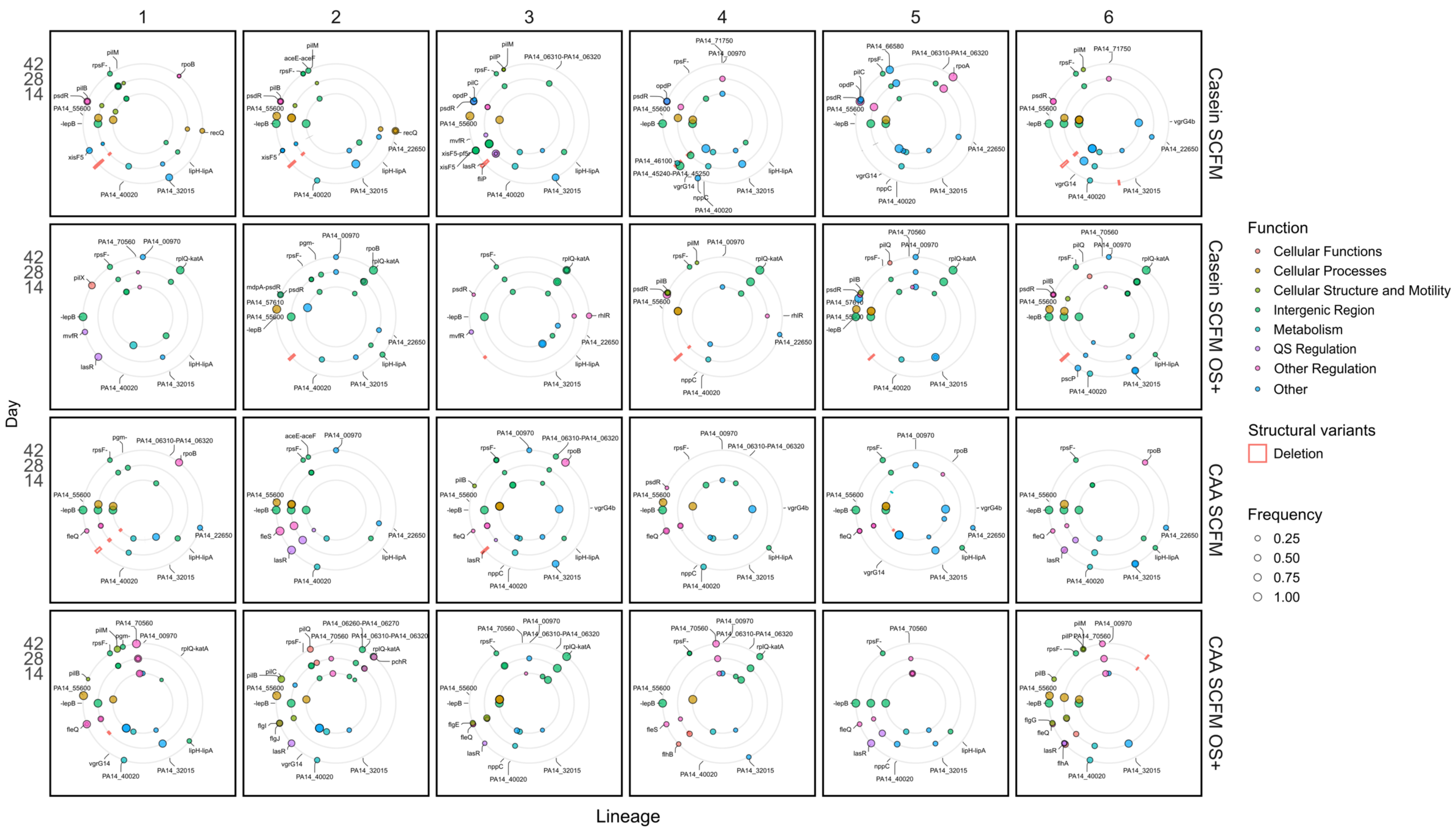

1
2
3
4
5
6
Casein SCFM
Casein SCFM OS+
CAA SCFM
CAA SCFM OS+
42
28
14
Day
Lineage
Function
Cellular Functions
Cellular Processes
Cellular Structure and Motility
Intergenic Region
Metabolism
QS Regulation
Other Regulation
Other
Structural variants
Deletion
Frequency
0.25
0.50
0.75
1.00

**Figure S5.** Mutational frequencies per evolving population over time. Each box represents a single population, with each row representing a selective environment (From top to bottom: Casein SCFM, Casein SCFM OS+, CAA SCFM, CAA SCFM OS+). Genes hit with mutations that reached high frequency (> 20%) at any detected sample are shown by their position on the chromosome (clockwise). The inner to outer rings show mutations on day 14, 28, and 42, respectively. Nonsynonymous SNPs and small indels are shown in filled circles with sizes ranging by frequency, coloured by gene functions (cellular functions in red, cell processes in orange, cellular structure and mobility in yellow-green, mutations in intergenic region in green, metabolism in cyan, QS regulation in purple, other regulation in pink, and other or unknown functions in blue). Deletions are shown in red boxes with height ranging by frequency, while no other structural variants were pronounced.

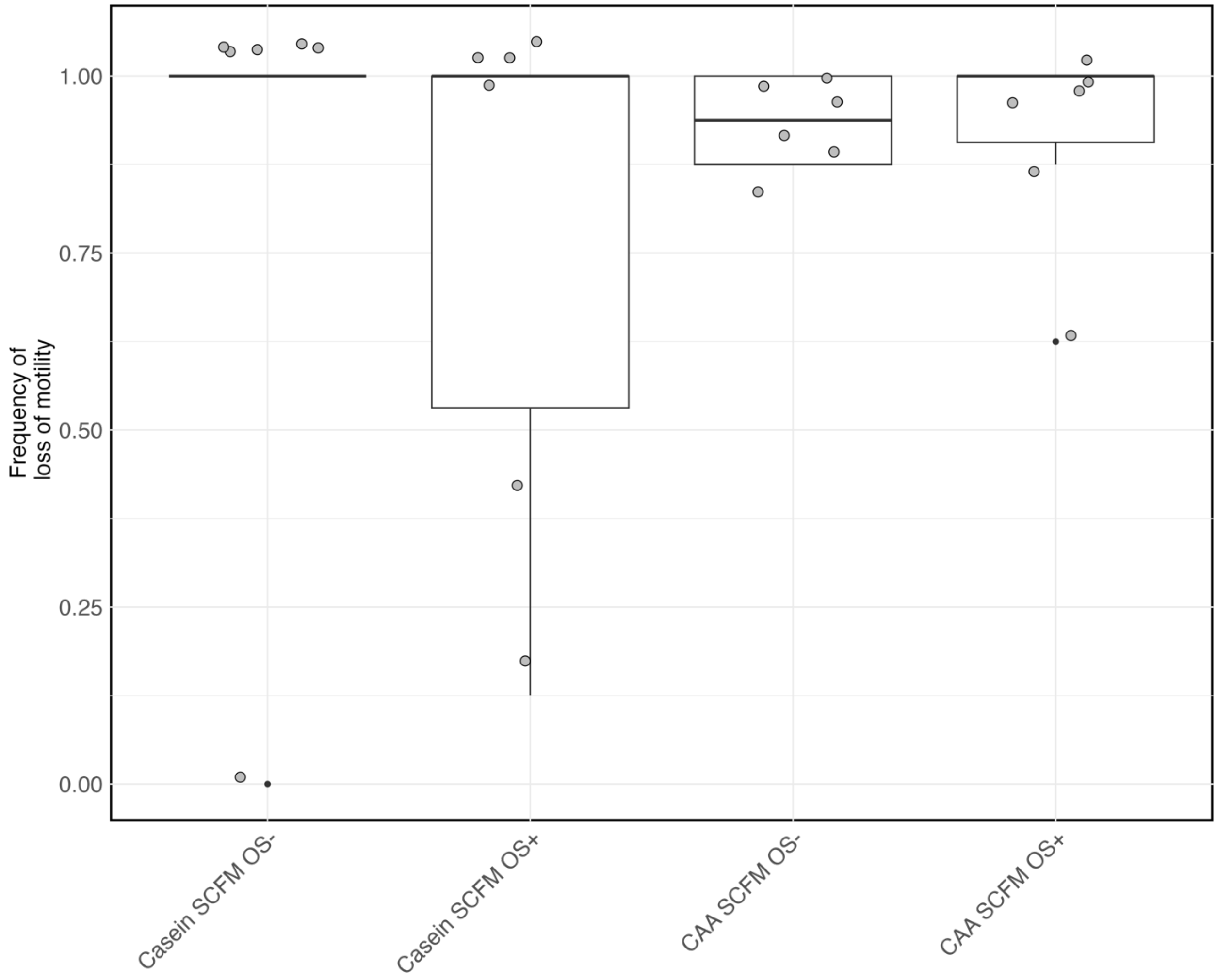


**Figure S6**. Frequencies of loss of flagellar swimming motility. Points show the frequency of clones (*n* = 8, per population) that have lost flagellar motility on day 42 per replicate population. Boxes show the mean and distribution per treatment.

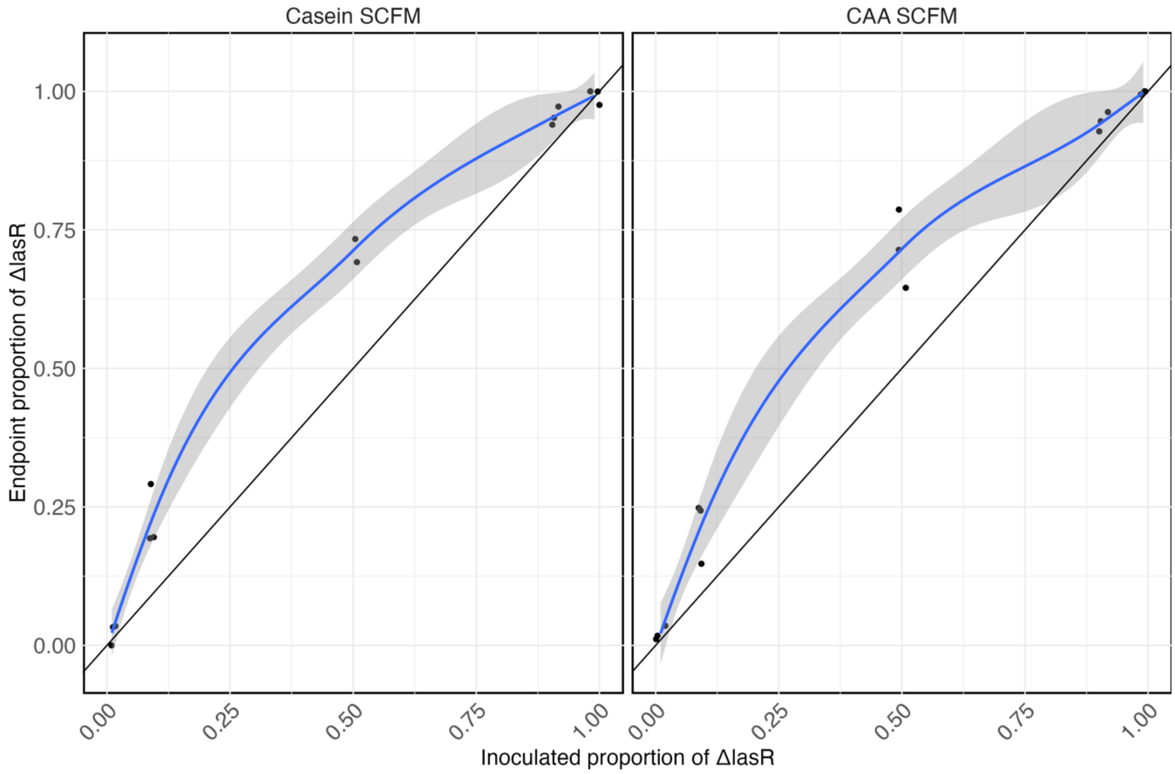


**Figure S7**. Competition assay between PA14 and PA14Δ*lasR* across starting ratios and nutrient conditions. Points show the endpoint proportion of PA14Δ*lasR* after 24 hours of co-culture plotted against the inoculated proportions (n = 3 biological replicates).

**Table S1.** Synonymous SNPs across all evolved lines (6 replicated populations per selective environments, 4 selective environments) and 3 timepoints (days 14, 28, and 42).

| Locus Label | Total Hits | Unique Hits | Locus Label | Total Hits | Unique Hits |
|---|---|---|---|---|---|
| *PA14_02560* | 3 | 1 | *PA14_61200* | 44 | 2 |
| *PA14_21020* | 1 | 1 | *PA14_65860* | 66 | 1 |
| *PA14_24780* | 17 | 1 | *argJ* | 2 | 1 |
| *PA14_31720* | 5 | 1 | *glgA* | 3 | 1 |
| *PA14_34270* | 3 | 3 | *mmsB* | 1 | 1 |
| *PA14_46100* | 1 | 1 | *nirB* | 6 | 1 |
| *PA14_47880* | 25 | 1 | *nppC* | 30 | 1 |
| *PA14_48890* | 1 | 1 | *rpoB* | 4 | 4 |
| *PA14_55600* | 8 | 3 | | | |

**Table S2.** Nonsynonymous SNPs and small indels across all evolved lines (6 replicated populations per selective environments, 4 selective environments) and 3 timepoints (days 14, 28, and 42).

| Locus Label | Total Hits | Unique Hits | Locus Label | Total Hits | Unique Hits |
|---|---|---|---|---|---|
| *PA14_00970* | 17 | 1 | *flgI* | 2 | 1 |
| *PA14_02560* | 5 | 1 | *flgJ* | 2 | 2 |
| *PA14_05740* | 1 | 1 | *flhA* | 2 | 1 |
| *PA14_08660* | 4 | 4 | *flhB* | 2 | 1 |
| *PA14_09380* | 5 | 3 | *fliI* | 2 | 1 |
| *PA14_10110* | 37 | 1 | *fliP* | 1 | 1 |
| *PA14_10260* | 1 | 1 | *fptA* | 2 | 1 |
| *PA14_13070* | 2 | 1 | *gabP* | 1 | 1 |
| *PA14_13150* | 1 | 1 | *hmgR* | 15 | 1 |
| *PA14_18720* | 2 | 1 | *lasR* | 15 | 8 |
| *PA14_19770* | 3 | 1 | *mdoH* | 15 | 1 |
| *PA14_20860* | 1 | 1 | *mdpA* | 2 | 1 |
| *PA14_21020* | 1 | 1 | *mexT* | 1 | 1 |
| *PA14_21120* | 2 | 1 | *mtnA* | 1 | 1 |
| *PA14_22650* | 11 | 1 | *mvfR* | 3 | 3 |
| *PA14_25620* | 1 | 1 | *nosD* | 26 | 1 |
| *PA14_26920* | 1 | 1 | *nppC* | 7 | 2 |
| *PA14_27550* | 2 | 1 | *opdP* | 3 | 3 |
| *PA14_31530* | 1 | 1 | *oprM* | 3 | 1 |
| *PA14_32015* | 36 | 3 | *orfN* | 5 | 1 |
| *PA14_32025* | 1 | 1 | *pcaB* | 4 | 1 |
| *PA14_32300* | 3 | 1 | *pchR* | 2 | 1 |
| *PA14_33150* | 2 | 1 | *pilB* | 11 | 8 |
| *PA14_33750* | 1 | 1 | *pilC* | 3 | 3 |
| *PA14_34270* | 1 | 1 | *pilE* | 1 | 1 |
| *PA14_35330* | 2 | 1 | *pilF* | 2 | 1 |
| *PA14_35770* | 3 | 2 | *pilM* | 7 | 6 |
| *PA14_35800* | 2 | 2 | *pilN* | 3 | 2 |

Continued on next page

Table S2 – continued from previous page

| Locus Label | Total Hits | Unique Hits | Locus Label | Total Hits | Unique Hits |
|---|---|---|---|---|---|

| | | | | | |
|---|---|---|---|---|---|
| *PA14_37360* | 3 | 1 | *pilO* | 1 | 1 |
| *PA14_40020* | 37 | 1 | *pilP* | 2 | 2 |
| *PA14_41340* | 2 | 2 | *pilQ* | 4 | 3 |
| *PA14_46020* | 1 | 1 | *pilR* | 2 | 2 |
| *PA14_46100* | 1 | 1 | *pilS* | 1 | 1 |
| *PA14_48010* | 4 | 1 | *pilW* | 1 | 1 |
| *PA14_51540* | 4 | 4 | *pilX* | 1 | 1 |
| *PA14_55600* | 46 | 3 | *pilY1* | 2 | 2 |
| *PA14_57610* | 2 | 1 | *pntB* | 1 | 1 |
| *PA14_62350* | 1 | 1 | *proB* | 47 | 1 |
| *PA14_62790* | 5 | 5 | *pscB* | 58 | 1 |
| *PA14_66580* | 2 | 1 | *pscP* | 3 | 1 |
| *PA14_68030* | 5 | 5 | *psdR* | 28 | 18 |
| *PA14_68150* | 3 | 3 | *pslI* | 11 | 1 |
| *PA14_70560* | 22 | 11 | *recQ* | 5 | 2 |
| *PA14_71100* | 11 | 1 | *rhlR* | 3 | 2 |
| *PA14_71740* | 1 | 1 | *rpoA* | 2 | 1 |
| *PA14_71750* | 2 | 2 | *rpoB* | 8 | 4 |
| *algC* | 60 | 1 | *trpI* | 1 | 1 |
| *argJ* | 21 | 1 | *vgrG14* | 11 | 2 |
| *fleQ* | 31 | 6 | *vgrG4b* | 4 | 1 |
| *fleS* | 4 | 2 | *xisF5* | 8 | 4 |
| *flgE* | 2 | 1 | *zbdP* | 1 | 1 |
| *flgG* | 2 | 1 | | | |

**Table S3.** Intergenic mutations across all evolved lines (6 replicated populations per selective environments, 4 selective environments) and 3 timepoints (days 14, 28, and 42).

| **Locus Label** | **Total Hits** | **Unique Hits** | **Locus Label** | **Total Hits** | **Unique Hits** |
|---|---|---|---|---|---|
| *-amtB* | 26 | 1 | *PA14_60920-PA14_60930* | 3 | 2 |
| *-hemH* | 1 | 1 | *PA14_68100-PA14_68110* | 1 | 1 |
| *-lepB* | 40 | 1 | *PA14_68630-PA14_68640* | 1 | 1 |
| *-metH* | 6 | 3 | *PA14_71260-PA14_71280* | 1 | 1 |
| *-plsB* | 11 | 3 | *aceE-aceF* | 2 | 1 |
| *-pncB2* | 8 | 1 | *acsB-* | 10 | 1 |
| *PA14_01660-PA14_01670* | 2 | 2 | *ansA-* | 9 | 2 |
| *PA14_04080-PA14_04090* | 1 | 1 | *dppA1-dppA2* | 3 | 1 |
| *PA14_05040-PA14_05050* | 14 | 1 | *exbD1-* | 1 | 1 |
| *PA14_06150-PA14_06160* | 3 | 3 | *ffh-rpsP* | 4 | 1 |
| *PA14_06260-PA14_06270* | 2 | 1 | *fimU-pilW* | 1 | 1 |
| *PA14_06310-PA14_06320* | 10 | 2 | *gabT-* | 2 | 1 |
| *PA14_12650-PA14_12670* | 24 | 1 | *ggt-ansB* | 1 | 1 |

Continued on next page

Table B.3 – continued from previous page

| **Locus Label** | **Total Hits** | **Unique Hits** | **Locus Label** | **Total Hits** | **Unique Hits** |
|---|---|---|---|---|---|

| | | | | | |
|---|---|---|---|---|---|
| *PA14_15540-PA14_15560* | 3 | 1 | *gyrA-serC* | 50 | 1 |
| *PA14_19230-PA14_19270* | 16 | 1 | *lipH-lipA* | 20 | 1 |
| *PA14_19290-PA14_19310* | 68 | 1 | *mdpA-psdR* | 1 | 1 |
| *PA14_21030-PA14_21040* | 1 | 1 | *napc-* | 38 | 1 |
| *PA14_21190-PA14_21210* | 3 | 1 | *nemO-* | 2 | 2 |
| *PA14_22860-PA14_22870* | 1 | 1 | *nuoD-nuoB* | 1 | 1 |
| *PA14_34410-PA14_34420* | 2 | 1 | *pgm-* | 3 | 1 |
| *PA14_35720-PA14_35730* | 2 | 2 | *rhlR-rhlI* | 2 | 2 |
| *PA14_35790-PA14_35800* | 1 | 1 | *rplQ-katA* | 32 | 3 |
| *PA14_45240-PA14_45250* | 2 | 1 | *rpoC-rpsL* | 2 | 2 |

Continued on next page

Table S3 – continued from previous page

| **Locus Label** | **Total Hits** | **Unique Hits** | **Locus Label** | **Total Hits** | **Unique Hits** |
|---|---|---|---|---|---|
| *PA14_45970-PA14_45980* | 4 | 2 | *rpsF-* | 60 | 4 |
| *PA14_46780-PA14_46800* | 21 | 1 | *sbrR-* | 12 | 2 |
| *PA14_48310-PA14_48320* | 1 | 1 | *sspA-* | 4 | 1 |
| *PA14_49030-PA14_49040* | 2 | 2 | *valS-* | 1 | 1 |
| *PA14_52920-PA14_52930* | 6 | 2 | *xisF5-pf5r* | 6 | 3 |

**Table S4.** Deletions across all evolved lines (6 replicated populations per selective environments, 4 selective environments) and 3 timepoints (days 14, 28, and 42).

| Locus Label | Total Hits | Locus Label | Total Hits |
|---|---|---|---|
| *pchB* | 2 | *PA14_45890* | 27 |
| *pchC* | 2 | *PA14_45910* | 27 |
| *pchD* | 2 | *PA14_45920* | 27 |
| *pchR* | 2 | *PA14_45930* | 27 |
| *pchE* | 2 | *lasI* | 27 |
| *pchF* | 2 | *lasR* | 27 |
| *PA14_34490* | 1 | *PA14_45970* | 27 |
| *PA14_34500* | 1 | *PA14_45980* | 25 |
| *PA14_34510* | 1 | *PA14_46010* | 19 |
| *PA14_34520* | 1 | *PA14_46020* | 17 |
| *PA14_34540* | 1 | *PA14_46030* | 15 |
| *PA14_34550* | 1 | *gbuR* | 14 |
| *PA14_34580* | 1 | *gbuA* | 14 |
| *PA14_34600* | 1 | *PA14_46080* | 14 |
| *gnuT* | 1 | *PA14_46100* | 13 |
| *PA14_34640* | 1 | *PA14_46110* | 9 |
| *gntR* | 1 | *PA14_46120* | 7 |
| *PA14_34670* | 1 | *PA14_46140* | 7 |
| *PA14_34680* | 1 | *PA14_46150* | 7 |
| *PA14_34690* | 1 | *PA14_46170* | 7 |
| *flhA* | 8 | *PA14_46180* | 7 |
| *PA14_45700* | 14 | *PA14_46200* | 5 |
| *PA14_45710* | 16 | *PA14_46220* | 2 |
| *flhB* | 17 | *aphA* | 2 |
| *fliR* | 20 | *PA14_46240* | 2 |
| *fliQ* | 20 | *PA14_46250* | 2 |
| *fliP* | 23 | *PA14_46260* | 2 |
| *fliO* | 25 | *PA14_46270* | 2 |
| *fliN* | 26 | *PA14_46290* | 2 |

Continued on next page

Table S4 – continued from previous page

| Locus Label | Total Hits | Locus Label | Total Hits |
|---|---|---|---|

| | | | |
|---|---|---|---|
| *fliM* | 27 | *PA14_46300* | 2 |
| *fliL* | 27 | *PA14_46310* | 2 |
| *PA14_45830* | 27 | *PA14_46320* | 2 |
| PA14_45840 | 27 | PA14_46330 | 2 |
| PA14_45850 | 27 | PA14_46360 | 2 |
| PA14_45870 | 27 | PA14_46370 | 1 |
| PA14_45880 | 27 | PA14_46400 | 1 |
| PA14_45840 | 27 | PA14_46330 | 2 |
| PA14_45850 | 27 | PA14_46360 | 2 |
| PA14_45870 | 27 | PA14_46370 | 1 |
| PA14_45880 | 27 | PA14_46400 | 1 |
